# Dataset factors associated with age-related changes in brain structure and function in neurodevelopmental conditions

**DOI:** 10.1101/2024.05.09.591294

**Authors:** Marlee M. Vandewouw, Yifan (Julia) Ye, Jennifer Crosbie, Russell J. Schachar, Alana Iaboni, Stelios Georgiades, Robert Nicolson, Elizabeth Kelley, Muhammad Ayub, Jessica Jones, Paul D. Arnold, Margot J. Taylor, Jason P. Lerch, Evdokia Anagnostou, Azadeh Kushki

**Affiliations:** Autism Research Centre, Bloorview Research Institute, Holland Bloorview Kids Rehabilitation Hospital, Toronto, ON, Canada; Institute of Biomedical Engineering, University of Toronto, Toronto, Canada; Division of Engineering Science, University of Toronto, Toronto, Canada; Department of Psychiatry, University of Toronto, Toronto, Canada; Department of Psychiatry, The Hospital for Sick Children, Toronto, Ontario, Canada; Department of Psychiatry and Behavioural Neurosciences, McMaster University, Hamilton, Canada; Department of Psychiatry, Western University, London, Canada; Department of Psychology, Queen’s University, Kingston, Canada; Centre for Neuroscience Studies, Queen’s University, Kingston, Canada; Department of Psychiatry, Queen’s University, Kingston, Canada; Division of Psychiatry, University of College London, London, United Kingdom; The Mathison Centre for Mental Health Research & Education, Cumming School of Medicine, University of Calgary, Calgary, Canada; Department of Diagnostic and Interventional Radiology, The Hospital for Sick Children, Toronto, Canada; Program in Neurosciences & Mental Health, The Hospital for Sick Children, Toronto, Canada; Department of Psychology, University of Toronto, Toronto, Canada; Department of Medical Imaging, University of Toronto, Toronto, Canada; Wellcome Centre for Integrative Neuroimaging, FMRIB, Nuffield Department of Clinical Neurosciences, University of Oxford, Oxford, United Kingdom; Department of Medical Biophysics, University of Toronto, Toronto, Canada; Institute of Medical Science, University of Toronto, Toronto, Canada

## Abstract

With brain structure and function undergoing complex changes throughout childhood and adolescence, age is a critical consideration in neuroimaging studies, particularly for those of individuals with neurodevelopmental conditions. However, despite the increasing use of large, consortium-based datasets to examine brain structure and function in neurotypical and neurodivergent populations, it is unclear whether age-related changes are consistent between datasets, and whether inconsistencies related to differences in sample characteristics, such as demographics and phenotypic features, exist. To address this, we built models of age-related changes of brain structure (regional cortical thickness and regional surface area; *N*=1,218) and function (resting-state functional connectivity strength; *N*=1,254) in two neurodiverse datasets: the Province of Ontario Neurodevelopmental network (POND) and the Healthy Brain Network (HBN). We examined whether deviations from these models differed between the datasets, and explored whether these deviations were associated with demographic and clinical variables. We found significant differences between the two datasets for measures of cortical surface area and functional connectivity strength throughout the brain. For regional measures of cortical surface area, the patterns of differences were associated with race/ethnicity, while for functional connectivity strength, positive associations were observed with head motion. Our findings highlight that patterns of age-related changes in the brain may be influenced by demographic and phenotypic characteristics, and thus future studies should consider these when examining or controlling for age effects in analyses.

## Introduction

The brain’s structure and function undergo complex and protracted changes throughout childhood and adolescence due to neurobiological processes such as synaptogenesis, myelination, and synaptic pruning (Grayson & Fair, 2017; Mills & Tamnes, 2020; Norbom et al., 2021). Neurodevelopmental differences, such as those associated with autism spectrum disorder (autism), attention-deficit/hyperactivity disorder (ADHD), and obsessive-compulsive disorder (OCD), can further impact the presentation of age-related changes, particularly given the heterogeneity in etiology, biology, and behaviour (Bruin et al., 2020; Lombardo et al., 2019; Wolfers et al., 2019). As such, age is a critical consideration in neuroimaging studies of neurotypical and neurodivergent populations.

While many studies directly examine age-related effects, age is often treated as a nuisance covariate (Hyatt et al., 2020). With increasing awareness of the importance of replicability in neuroimaging studies (Poldrack et al., 2017), it is becoming more common to leverage multiple consortium-based datasets to examine brain structure and function in neurotypical and neurodivergent populations (Abrol et al., 2023; Grotzinger et al., 2023; Marek et al., 2022; Nicolaisen-Sobesky et al., 2022; Romer et al., 2019; Vandewouw et al., 2023). However, these datasets differ in composition across factors which may influence age-related effects, such as participants’ demographics (e.g., age, sex and gender, race and ethnicity, socioeconomic status), diagnosis, co-occurring conditions, and other phenotypic variables. As a result, it is possible that the patterns of age-related changes in brain structure and function differ between datasets. Since it is common to apply the same assumptions about the development of neurobiology across datasets (e.g., linearity), the presence of dataset-specific developmental effects would be problematic for downstream analyses, challenging replicability.

Studies have examined the consistency of age-related changes in brain structure and function across neurotypical cohorts (Cao et al., 2017; Herting et al., 2018; Madan & Kensinger, 2017; Mills et al., 2016; Morandini et al., 2021; Tamnes et al., 2017; Wang et al., 2021). For example, yearly percentage changes in cortical thickness and surface area across childhood and adulthood have been shown to differ between four samples of typically developing individuals, with the precise patterns varying across lobes of the brain (Tamnes et al., 2017). White matter volume has also been shown to peak at different age ranges in different cohorts, with those from the United States showing decreased volume across age compared to European samples (Mills et al., 2016).

These findings have important implications for those studies which are not only directly examining age-related effects, but also those which treat age as a covariate, as controlling for age in the same manner does not consider these differences. Yet, no studies have examined whether these changes are consistent between cohorts of neurodivergent children, and if not, what factors may be contributing to the inconsistencies. This is particularly critical to investigate given the large variability in demographic and phenotypic characteristics of samples across different cohorts, even when the same diagnostic labels are studied.

While there is a wealth of evidence showing that demographic and clinical factors can impact brain development (e.g., neurodivergence), most of this work has taken a case-control approach, where differences from neurotypical trajectories are established at the “average” level. On the other hand, there is a considerable amount of developmental variation at the individual level (Foulkes & Blakemore, 2018), with emerging evidence that clinical features of autism and ADHD are associated with individual deviations from neurotypical maturational trajectories (Bethlehem et al., 2020; Floris et al., 2021; Kessler et al., 2016; Looden et al., 2022; Marquand et al., 2016; Oliveira-Saraiva & Ferreira, 2023; Parkes et al., 2021; Tunç et al., 2019; Wolfers et al., 2019; Zabihi et al., 2019). Thus, to understand why datasets may exhibit distinct age-related changes of neurobiology, it is important to consider how demographic and clinical factors impact individual variations.

To this end, we examined age-related changes of brain structure (regional cortical thickness and surface area) and function (functional connectivity strength) in two transdiagnostic datasets of neurodiverse children and adolescents, namely the Province of Ontario Neurodevelopmental network (POND) and the Healthy Brain Network (HBN). We first built dataset-specific developmental models and evaluated whether deviations from these models differed between the POND and HBN individuals, separately for males and females. Next, we explored whether the patterns of between-dataset differences were associated with demographic and phenotypic variables. We have previously reported that POND and HBN differ in their phenotypic profiles, specifically full-scale intelligence quotient (FSIQ), social communication difficulties, and ADHD symptoms (Vandewouw et al., 2023). Thus, we hypothesize that deviations from age-related models of brain structure and function will differ between these datasets, and the deviations will be associated with these phenotypic measures.

## Methods

### Participants

Neuroimaging and phenotype data were obtained from two independently collected datasets: the Province of Ontario Neurodevelopmental network (POND; exported April 2023), and the Healthy Brain Network (HBN; Release 9). For both datasets, participants who were between 5-19 years of age and who were either neurotypical or had diagnoses of autism, attention-deficit/hyperactivity disorder (ADHD), or obsessive-compulsive disorder (OCD) were selected. Details on the diagnostic assessments are provided elsewhere (Vandewouw et al., 2023). Structural magnetic resonance imaging (sMRI) data were available for 868 POND (255 ADHD, 377 autism, 75 OCD, 161 TD) and 1,505 HBN (1,100 ADHD, 162 autism, 36 OCD, 207 TD) participants and resting-state functional magnetic imaging (rs-fMRI) data were available for 864 POND (211 ADHD, 385 autism, 92 OCD, 185 TD) and 1,459 HBN (1,068 ADHD, 160 autism, 29 OCD, 202 TD) participants. Informed consent and/or assent was obtained from the participants and/or caregivers. The current study was approved by the institution’s research ethics board, and both consortia studies were approved by the appropriate boards.

### Data acquisition

For both datasets, T1-weighted magnetization-prepared rapid gradient echo (MPRAGE) sequences (sMRI data) and five minutes of resting-state data with an echo planar imaging (EPI) sequence (rs-fMRI data) were acquired. In POND, the rs-fMRI data were obtained as participants watched either a movie of their choosing or the naturalistic movie paradigm akin to a screensaver, *Inscapes* (Vanderwal et al., 2015), while HBN participants viewed a fixation cross. While HBN collected two five-minute resting-state acquisitions, only one was selected to be consistent with POND. Acquisition parameters for the sMRI and rs-fMRI protocols for each POND and HBN site are presented in **Supplemental Table 1**.

Biological sex was collected as part of both studies and used in the current analysis; insufficient data on gender was available at the time of export. Phenotypic measures of core and co-occurring differences for the diagnostic categories (see **Supplemental Table 2**), socioeconomic status, and race and ethnicity data were also available for both datasets and were used to characterize each sample. Socioeconomic status was determined using the caregiver’s highest level of education and annual household income. Race and ethnicity data were collected for both datasets according to each country’s census guidelines and were categorized into five racial/ethnic categories: Hispanic/Latino, non-Hispanic Asian, non-Hispanic Black, non-Hispanic White, and Multi-Racial/Other (Cardenas-Iniguez & Gonzalez, 2024).

### Image processing

The sMRI data were processed using the CIVET pipeline (version 2.1.0; (Ad-Dab’bagh et al., 2006)) on CBRAIN (Sherif et al., 2014). CIVET first registers the T1-weighted image to stereotaxic space (Collins et al., 1994) using the Montreal Neurological Institute template (MNI ICBM152; (Fonov et al., 2009)) and corrected for non-uniformities (Sled et al., 1998). A brain mask was extracted (Smith, 2002), and tissues were classified into white matter, gray matter, and cerebrospinal fluid using a discrete tag point classification and a correction for partial volumes (Tohka et al., 2004; Zijdenbos et al., 1998). The white matter and pial surfaces (40,962 vertices per hemisphere) were extracted using the marching-cubes algorithm (Kabani et al., 2001; Kim et al., 2005; MacDonald et al., 2000). Vertex-wise cortical thickness was calculated as the Euclidean distance between reference vertices of the white and pial surfaces (Ad-Dab’bagh et al., 2005; Lerch & Evans, 2005). Vertex-wise cortical surface area and volumes were calculated as local variations of contraction and expansion relative to the distribution of vertices (Lyttelton et al., 2009). Surface-based diffusion smoothing kernels were applied (thickness: 30mm full-width at half-maximum (FWHM); area and volume: 40mm FWHM) (Boucher et al., 2009). Regional measures of cortical thickness and surface area were extracted for 76 brain regions defined using the Automated Anatomical Labelling (AAL) atlas (Tzourio-Mazoyer et al., 2002) by averaging across the vertices per region. The AAL atlas was chosen to be consistent with most studies using the CIVET pipeline, including work from our group (Kushki et al., 2019, 2021; Sadat-Nejad et al., 2023; Sanjeevan et al., 2020; Sussman et al., 2015). The CIVET automatic quality control pipeline was used, details of which are described elsewhere (Hammill et al., 2021; Kushki et al., 2019, 2021). Of the remaining participants that passed quality control, propensity score matching (Rosenbaum & Rubin, 1985) was used to identify a smaller subset of participants to ensure the datasets showed no significant difference in median age or male-to-female ratio (see (Vandewouw et al., 2023), for further details). Across both datasets, ComBat harmonization (Fortin et al., 2018; Johnson et al., 2007) was used to adjust all measures for site effects, followed by the regression of mean cortical thickness and total surface area for the regional measures of thickness and area, respectively. For both steps, an iterative sampling procedure was performed to increase robustness (10,000 iterations, each selecting 63.2% of the sample (Da-ano et al., 2020)).

The rs-fMRI data preprocessing was performed using fMRIPrep (Esteban et al., 2019, 2020), and is described in detail elsewhere (Vandewouw et al., 2023). As part of the pipeline, measures of framewise displacement (FD; (Power et al., 2014)) and the standardized derivative of root mean square variance over voxels (DVARS; (Power et al., 2014)) were obtained. Participants were required to have at least 2/3 of their data within the recommended motion thresholds (FD: 0.5mm; DVARS: 1.5) to pass quality control. Of the remaining participants, propensity score matching was used to identify a smaller subset of individuals for which there was not a statistically significant between-dataset differences in median age, male-to-female ratio, and median head motion (Vandewouw et al., 2023). The Schaefer cortical (Schaefer et al., 2018) and Melbourne subcortical (Tian et al., 2020) atlases were used to define 232 brain regions, which have been shown to produce functional network topology that is more similar across network construction pipelines compared to the AAL and other alternative functional parcellations (Luppi & Stamatakis, 2021). Pairwise Pearson correlations were computed between region-averaged time series to construct rs-fMRI connectomes. ComBat harmonization (Da-ano et al., 2020; Fortin et al., 2018; Johnson et al., 2007) was used to adjust the connectivity values for site effects across both datasets (10,000 iterations, each selecting 63.2% of the sample). The resulting connectomes were thresholded with orthogonal minimum spanning trees to remove spurious connections (Dimitriadis et al., 2017), and the functional connectivity strength of each brain region was extracted.

### Modeling age-related changes

Models of age-related changes in the regional sMRI (area and thickness) and rs-fMRI (connectivity strength) measures were built using Bayesian linear regression (BLR; (Huertas et al., 2017)). Unlike traditional regression, BLR estimates the posterior probability of the regression coefficient, capturing the expected variation or uncertainty around the predicted age-related change; this enables the computation of a statistical *z*-score, or deviation score, that is similar to a residual but also considers the uncertainty around the prediction (Huertas et al., 2017; Rutherford et al., 2022). BLR is comparable but less computationally expensive than similar techniques such as Gaussian process regression (Fraza et al., 2021; Huertas et al., 2017). We used the BLR implementation from the Predictive Clinical Neuroscience toolkit (version 0.25; https://pcntoolkit.readthedocs.io/en/latest) in Python (version 3.9), applying a B-spline basis expansion of age with three evenly spaced knots to model non-linearities, using warping to model normal and non-normal distributions in the brain measures, and using the Powell method for optimization of the Bayesian hyper-parameters.

Dataset (POND and HBN; *N*=2) and sex (males and females; *N*=2) specific models were built for each measure (regional cortical thickness, regional cortical surface area, and functional connectivity strength; *N*=3), resulting in 12 models. To promote generalizability, 80% of participants of a given dataset were used to build a model, with 20% withheld for testing. This procedure was repeated 10,000 times, each time with a different randomly selected subset of participants. To describe the performance of the models, the model fit was evaluated by examining the mean standardized log-loss (MSLL; (Rasmussen & Williams, 2006)), which quantifies the error between the predicted and true mean of the data while also accounting for the predictive variances, and has an upper bound of 0, with negative values indicating a better model fit (Rutherford et al., 2022). Deviation scores (*z*-scores) were calculated from a built model under two test scenarios: the model built on 80% of a given dataset was tested on 1) the remaining 20% of participants from the same dataset, and 2) the participants from the other dataset. **Figure 1** depicts an example of this process where the model is built using 80% of the participants in POND and tested on the remaining 20% of participants from POND, and 100% of the HBN participants. For robustness, the analysis was also performed with a 70%/30% train/test split. Median model fit metrics and deviation scores were computed across all iterations. This procedure was performed separately for males and females, with both the full samples and smaller, matched samples.

**Figure 1:**
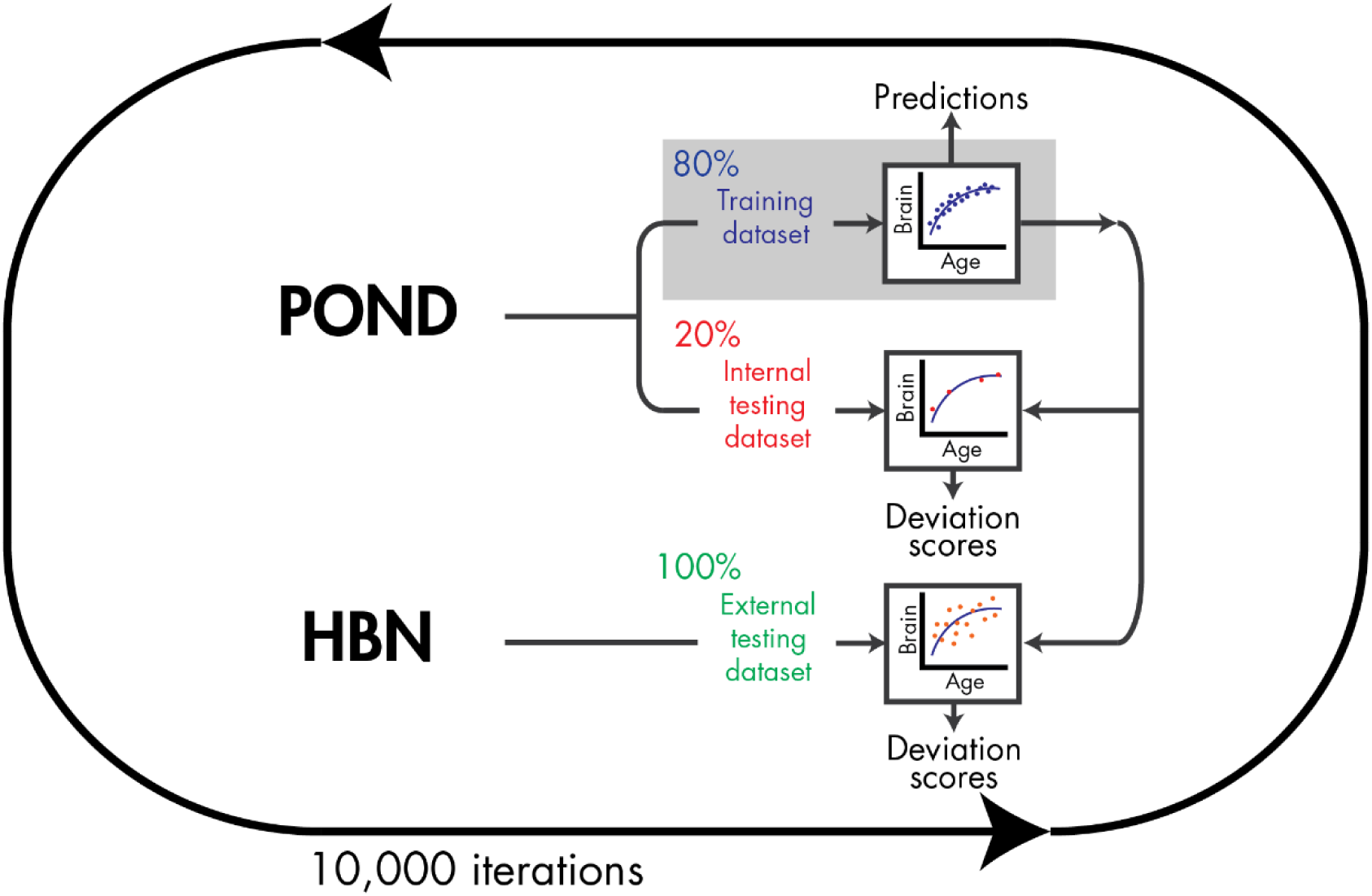
An example of the approach for constructing comparing the dataset- and sex-specific BLR models.

### Statistics

First, differences between the two datasets in their demographic and clinical characteristics were examined, separately for males and females. Chi-squared tests, Kruskal-Wallis tests (given non-normality according to Shapiro-Wilks tests), and ordinal regressions were used for categorical, continuous, and ordinal variables, respectively, reporting Cramer’s *v*, eta-squared, and odds ratio effect sizes; in all cases, significance was held at p<0.05.

For each of the brain measures, we compared the deviations between the datasets; this was performed with both POND and HBN used as the training dataset, and separately for males and females. One-way ANOVAs or Kruskal-Wallis tests were used where appropriate, resulting *p*-values were adjusted using the false discovery rate (FDR) method, holding significance at *q*<0.05 and reporting eta-squared effect sizes.

Given differences in demographic (socioeconomic status and race/ethnicity) and clinical (diagnosis and phenotypic measures) variables between POND and HBN, we explored what factors may be contributing to any identified between-dataset differences. To do so, for each brain measure, we extracted the deviations for each participant from their dataset- and sex-specific model and used generalized linear mixed-effects models to examine their associations with these variables; dataset and sex were modeled as random factors. Resulting *p*-values were FDR-corrected across the brain regions, and significance was held at *q* < 0.05.

## Results

### Participants

For the sMRI dataset, 754 HBN (214 female, 540 male) and 825 POND (270 female, 555 male) children and adolescents were included in the analysis. Given the difference in age between the two datasets (females: *H*(1)=104.35, *p*<.001, *η*^2^=.04; males: *H*(1)=203.75, *p*<.001, *η*^2^=.03), a smaller, matched subset of participants was identified for which no significant difference remained, with each dataset consisting of 609 (178 females, 431 males) individuals. Participant demographics for the matched sample are presented in **Table 1**, with phenotypic measures, socioeconomic status, and race/ethnicity for each dataset are presented in **Table 2**. Co-occurring conditions are summarized in **Supplemental Table 3**.

**Table 1:** Participant demographics for the sMRI sample.

|  |  | HBN | POND | Test-Statistic<sup>1</sup> | p-value | Effect size |
| --- | --- | --- | --- | --- | --- | --- |
| **Females** | **N** | 178 | 178 | – | – | – |
|  | **ADHD** | 124 | 51 | 76.33 | <.001 | .46 |
|  | **Autism** | 7 | 56 |  |  |  |
|  | **OCD** | 9 | 24 |  |  |  |
|  | **Dx (N) TD** | 38 | 47 |  |  |  |
|  | **Age<sup>2</sup> (years)** | 10.1 [8.5, 13.3] | 11.1 [9.0, 13.3] | 2.61 | .107 | <.01 |
| **Males** | **N** | 431 | 431 | – | – | – |
|  | **ADHD** | 304 | 145 | 141.73 | <.001 | .41 |
|  | **Autism** | 55 | 190 |  |  |  |
|  | **OCD** | 11 | 33 |  |  |  |
|  | **TD** | 61 | 63 |  |  |  |
|  | **Age<sup>2</sup> (years)** | 10.6 [8.6, 13.0] | 10.9 [9.0, 13.0] | 2.07 | .150 | <.01 |

HBN: Healthy Brain Network; POND: Province of Ontario Neurodevelopmental network; ADHD: attention-deficit/hyperactivity disorder; OCD: obsessive-compulsive disorder; TD: typically developing; <sup>1</sup>Kruskal-Wallis *H*-statistic for age with eta-squared effect size, and chi-squared statistic for diagnosis with Cramer's *v* effect size; <sup>2</sup>Age is reported as median [IQR].

**Table 2:**
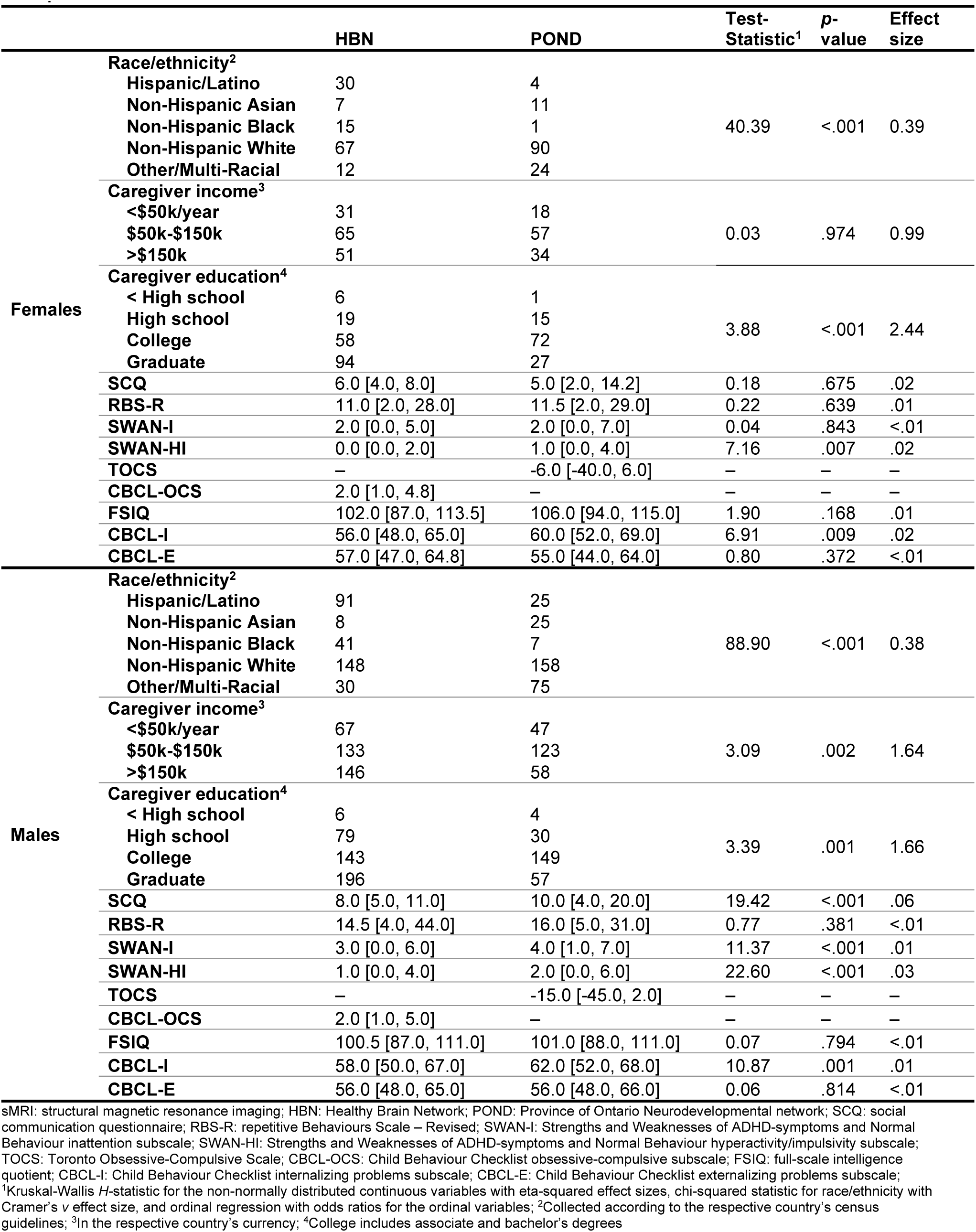
Socioeconomic status, race/ethnicity, and phenotypic measures for the sMRI sample.

For the fMRI dataset, 1072 HBN (349 females, 723 males) and 720 POND (207 females, 513 males) participants were included in the analysis. Significant between-dataset differences in age (females: *H*(1)=21.34, *p*<.001, *η*^2^=.04; males: *H*(1)=40.82, *p*<.001, *η*^2^=.03) and motion (females: *H*(1)=36.50, *p*<.001, *η*^2^=.03; males: *H*(1)=29.50, *p*<.001, *η*^2^=.01) were observed. Thus, a smaller, matched subset of children and adolescents with no such differences was identified, with each dataset consisting of 627 (189 female, 438 male) individuals (**Tables 3** and **4**). Co-occurring conditions are summarized in **Supplemental Table 4**. In this matched sample, there were no significant between-diagnosis differences in head motion in the POND (*H*(3)=0.51, *p*=.676, *η*^2^=0.01) nor HBN (*H*(3)=1.07, *p*=.362, *η*^2^=0.02) females. In the males, diagnosis was marginally significant in POND (*H*(3)=2.68, *p*=.046, *η*^2^=0.02) but not significant in HBN (*H*(3)=1.79, *p*=.149, *η*^2^=0.01).

**Table 3:**
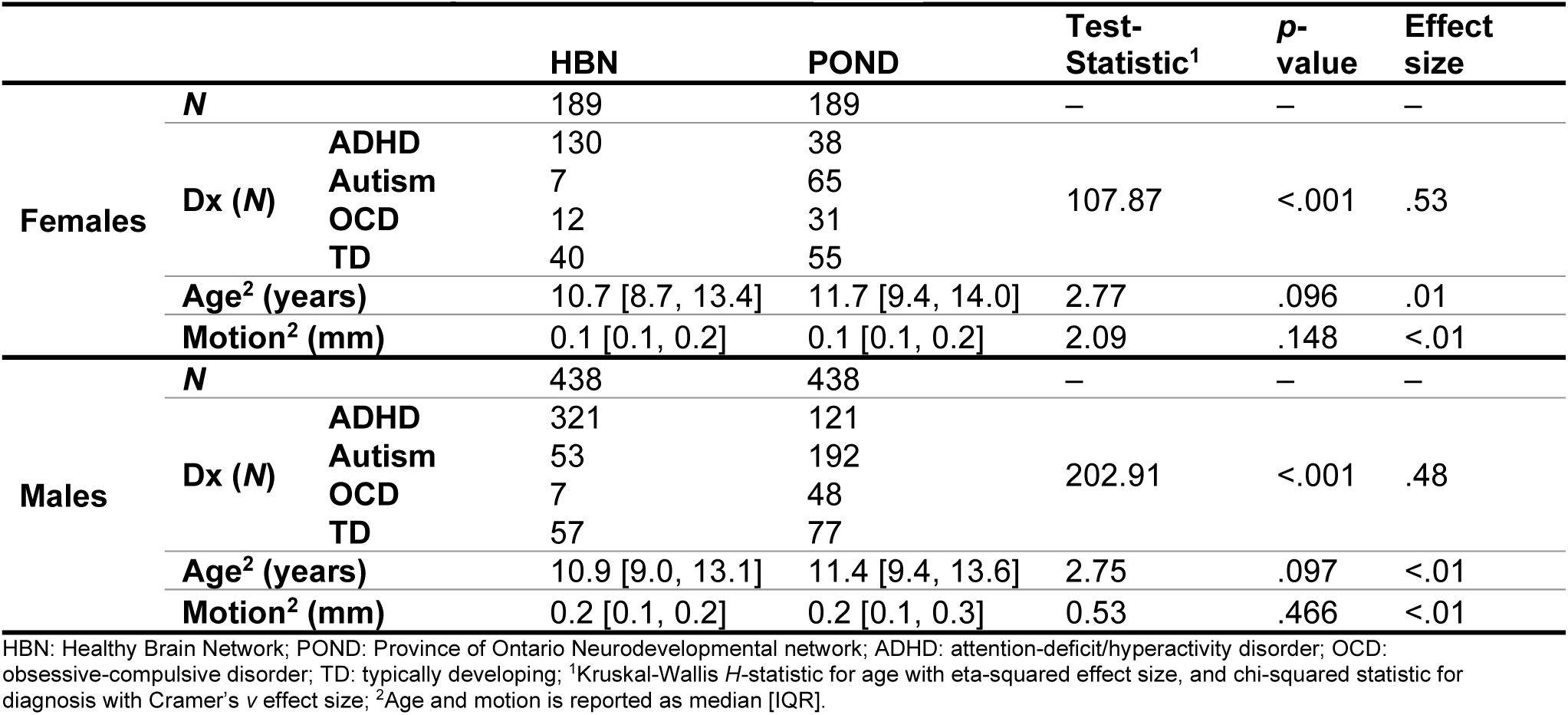
Participant demographics for the fMRI sample.

**Table 4:**
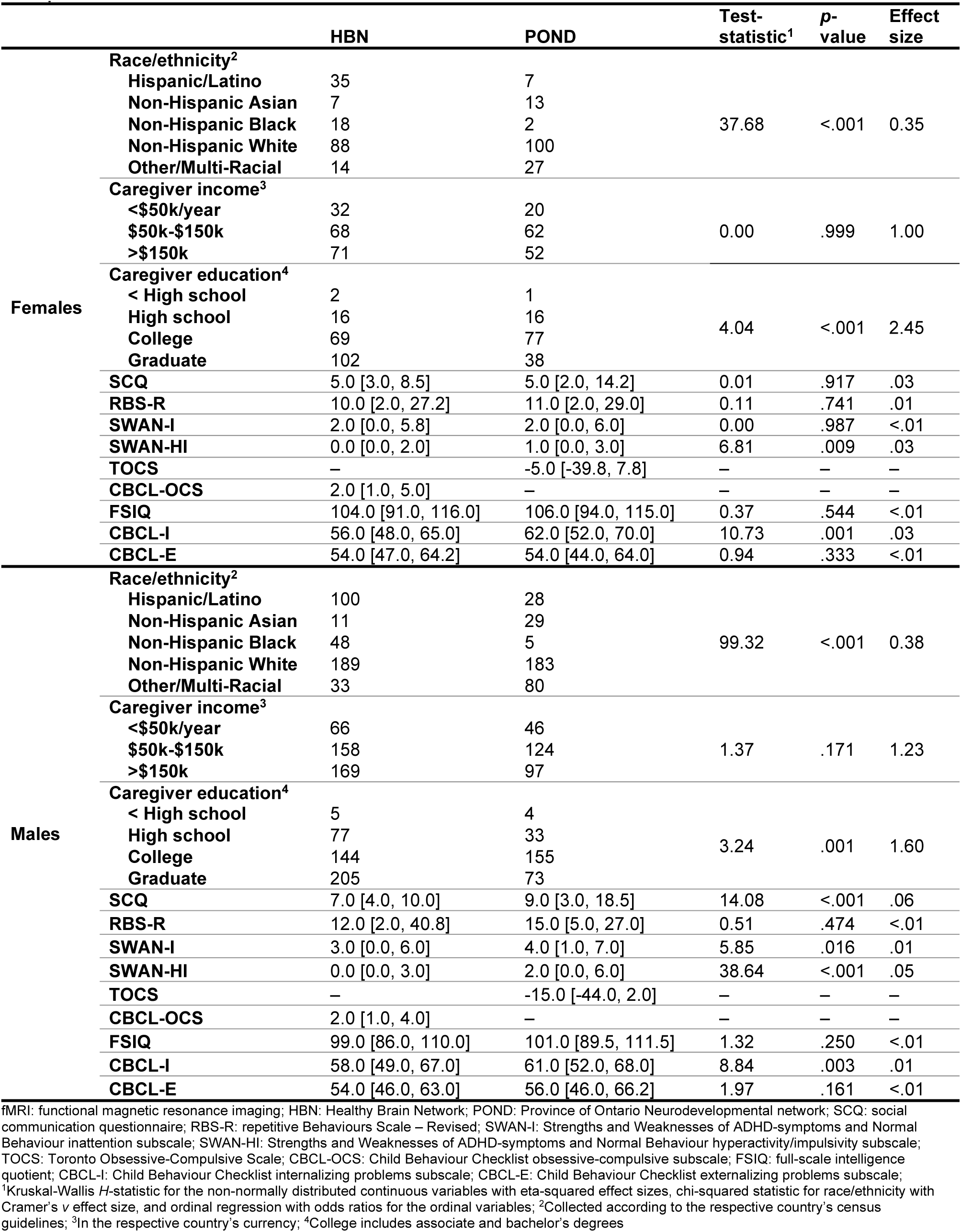
Socioeconomic status, race/ethnicity, and phenotypic measures for the fMRI sample.

### Dataset differences in deviations

Models of age-related changes in cortical thickness, surface area, and functional connectivity strength were fit separately for each dataset and sex. To evaluate the model performance, the distributions of MSLL across all brain regions are presented in **Supplemental Figure 1**, and distributions of deviation scores are presented in **Supplemental Figure 2**. The deviation scores for the age models were compared between datasets, separately for the males and females. Significant differences were observed for cortical surface area and functional connectivity, but not cortical thickness, with significant differences presented in **Figure 2**. Findings were robust to the size of the training and test sets (**Supplemental Figure 3**). Significant between-dataset differences within each diagnostic label are presented in **Supplemental Figure 4**. To explore what factors may be contributing to the observed differences, associations between deviations and the demographic variables (age, sex, race/ethnicity, caregiver highest level of education, and annual household income) and phenotypic variables (social communication difficulties, repetitive behaviours, inattention, hyperactivity/impulsivity, obsessive-compulsive traits, intelligence, internalizing problems, and externalizing problems) were investigated.

**Figure 2:**
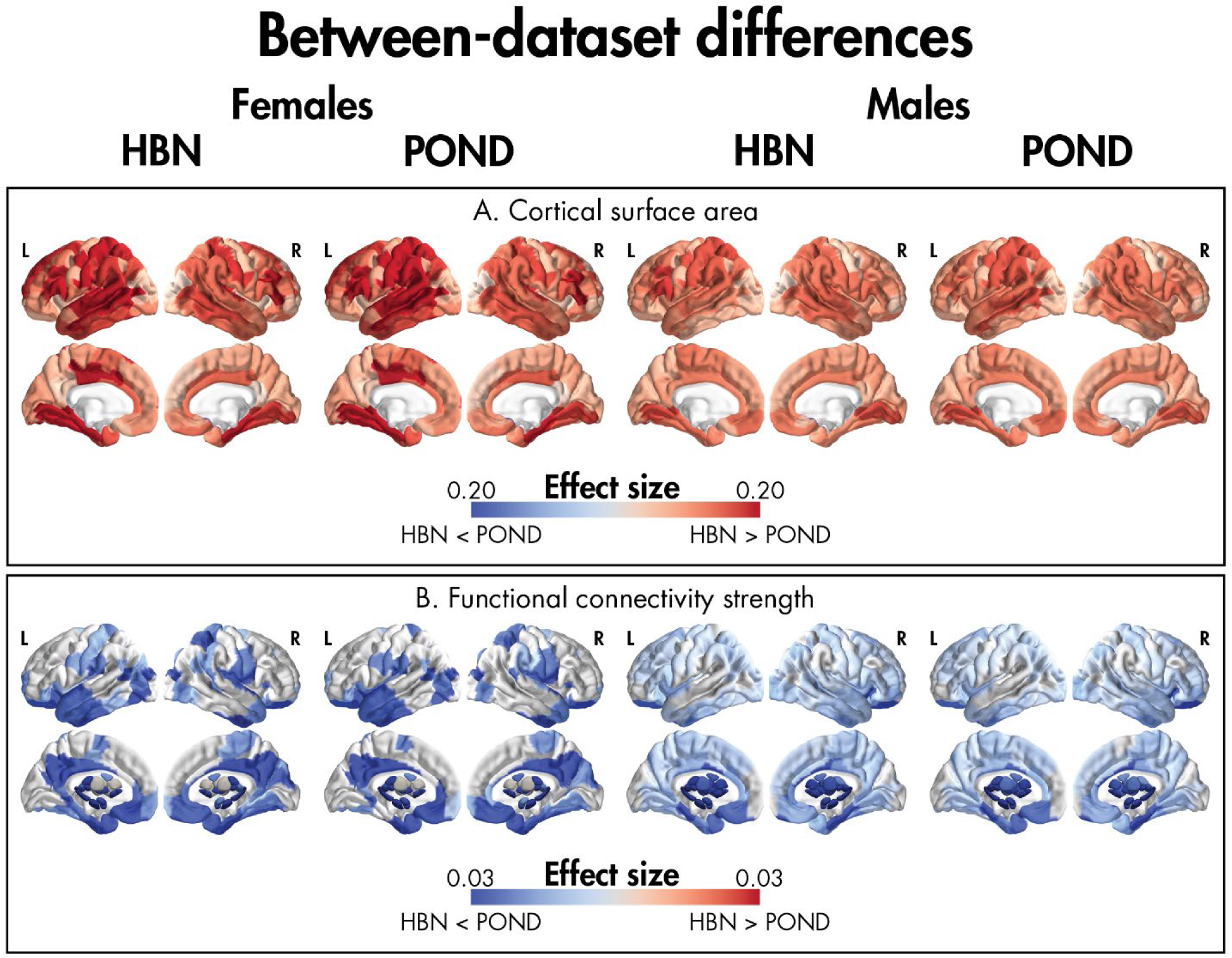
Effect sizes (eta-squared) for significant between-dataset differences in deviations from age-related models of cortical surface area (A) and functional connectivity strength (B) for the females (left) and males (right) when the models were built using each dataset (HBN: Healthy Brain Network, POND: Province of Ontario Neurodevelopmental network). No differences in cortical thickness were observed.

For regional measures of cortical surface area (**Figure 2A**), deviation scores differed between the datasets throughout the brain. When examining the median deviation scores for the significant regions, deviations for the POND participants were negative when the models were built using the HBN participants, indicating that the HBN models overestimated surface area for the POND individuals; on the other hand, POND models underestimated surface area in the HBN participants (positive deviation scores). Significant associations between deviations and race/ethnicity were observed in the left orbital part of the middle frontal gyrus, left medial superior frontal gyrus, right supplementary motor area, right angular gyrus, right middle and inferior occipital gyri, right fusiform gyrus, right parahippocampal gyrus, and right median cingulate and paracingulate gyrus (**Supplemental Figure 5A**). Post-hoc tests indicated that while deviations in White participants were closer to zero, deviations in minoritized participants were increased in magnitude. All other associations were insignificant. For regional measures of cortical thickness, no regions showed significant between-dataset differences in the deviations.

For functional connectivity strength (**Figure 2B**), significant differences in the deviation scores between POND and HBN were observed throughout the brain, with more widespread effects observed in the males (HBN: 202 regions, POND: 199 regions), albeit with reduced overall effect sizes, compared to females (HBN: 135 regions, POND: 122 regions). When models were built using the HBN individuals, median deviation scores indicated that they underestimated functional connectivity strength in POND; conversely, POND models overestimated connectivity strength in the HBN participants. Deviation scores were significantly positively correlated with head motion (**Supplemental Figure 5B**).

## Discussion

In this study, using data from two consortia, namely POND and HBN, we built dataset-specific models that describe age-related changes in measures of brain structure and function. We determined that deviations from the models were significantly different between the two datasets for measures of regional cortical surface area and functional connectivity strength throughout the brain, but not regional cortical thickness; these differences were observed within both males and females. For cortical surface area, the patterns of differences were associated with race/ethnicity, suggesting that it may be contributing to the between-dataset findings. For functional connectivity strength, positive associations were observed with head motion.

Overall, our findings highlight that patterns of age-related changes in the brain may not be consistent between cohorts of neurodiverse individuals, and these inconsistencies may be related to differences in the demographic and phenotypic composition of the sample. While previous work has identified factors such as methodology and medication (Bednarz & Kana, 2018) as contributors to the lack of convergence in the literature on the nature and directionality of brain differences in neurodevelopmental conditions, we hypothesize that differences in age-related changes between cohorts, potentially driven by differences in measures like race/ethnicity and head motion, are also contributing to the lack of consistent findings. Even though age may not be the focus of a particular study, adjusting for age in the same manner in multiple cohorts despite the presence of cohort-specific age-related effects can produce downstream effects in subsequent analyses.

For the regional measures of cortical surface area, differences in deviations from age models differed between POND and HBN in every brain region, irrespective of which group was used to construct the model. The magnitude of this effect differed throughout the brain, for example, larger effects were observed in parietal regions. The development of cortical surface area shows the largest changes across childhood, adolescents, and early adulthood in the parietal lobe (Tamnes et al., 2017). Thus, we hypothesize that while dataset-specific age-related changes of cortical surface area are pervasive, brain regions which show more drastic developmental differences may show more exaggerated effects.

Deviations from age models of cortical surface area were also larger in magnitude for minoritized compared to White individuals, including frontal, occipital, and cingulate regions. Given that deviations from the models were higher in magnitude in the minoritized compared to white participants, we hypothesize that this may reflect the effect of underrepresentation of minoritized individuals in the two datasets. Interestingly, despite race/ethnicity being significantly associated with measures of socioeconomic status (household income, highest education level), we did not observe an association between socioeconomic status and the deviations alongside race/ethnicity. Potential mechanisms contributing to racial/ethnic differences in brain structure have been suggested to include, for example, childhood adversity (Dumornay et al., 2023), although we did not explicitly test this in our study.

While between-dataset differences in deviations from age models were observed for regional cortical surface area, this finding did not extend to regional cortical thickness. The development of cortical thickness and surface area are driven by different genetic and biological processes and show distinct maturational trajectories (Geschwind & Rakic, 2013; Panizzon et al., 2009; Tamnes et al., 2017; Vijayakumar et al., 2016; Wierenga et al., 2014). Surface area has also been shown to uniquely relate to general psychopathology, while this association is not observed with cortical thickness in several well-powered studies (Jalbrzikowski et al., 2019; Mewton et al., 2022; Romer et al., 2023; Schmaal et al., 2016), although we acknowledge that smaller studies have reported a link between psychopathology and cortical thickness (e.g., (Luking et al., 2022)). Thus, given the differences between POND and HBN in the core and co-occurring domains of neurodevelopmental conditions, we hypothesize that the unique relationship between psychopathology with surface area is why differences in deviations are restricted to this structural measure rather than cortical thickness.

We also observed differences between POND and HBN in deviations from models describing changes in functional connectivity strength with age, with the largest effects observed in limbic, default mode, and the subcortical regions. Deviations from the models of age-related effects were also shown to be positively associated with head motion, with, again, the largest effect sizes in limbic, default, mode, and subcortical regions. Head motion is known to increase functional connectivity (Satterthwaite et al., 2013), and head motion is higher in HBN compared to POND, possibly due to the use of a fixation-cross resting-state (HBN) compared to movie paradigms (POND) (Vanderwal et al., 2015). Thus, together with the noticeable overlap in the spatial pattern of effects, we suggest that head motion may be driving deviations in age-related changes of functional connectivity. This conclusion is supported by the largest effect sizes being localized to the subcortical structures, which are located distally from other brain regions, and thus may be more impacted by head motion due to its effect on long-range connectivity (van Dijk et al., 2012). Note that while differences in head motion are often reported between children with and without neurodevelopmental conditions (Vandewouw et al., 2023), in our matched samples, this was not the case – only marginal differences were observed in the POND males, with no other groups showing this pattern. Coupled with the lack of associations between diagnosis and model deviations, it does not appear that the correlations between deviations and head motion were being driven by diagnosis.

A strength of this study is the stratification of the analyses by biological sex. Sex and gender differences in the prevalence rates of neurodevelopmental conditions have long contributed to biases in existing research samples and neuroimaging literature (Mo et al., 2021). Specifically, the male preponderance of many of these conditions has resulted in a poor understanding of females with these conditions (Mo et al., 2021). This has resulted in calls for stratifying by sex and gender in neurodevelopmental research (Bölte et al., 2023), which motivated the decision to fit sex-specific age-related models. However, we recognize that the binarized measure of biological sex used in this study is an oversimplification of sex and gender; future work should account for more nuanced identities.

The results of this study should be interpreted considering several limitations. This study is cross-sectional, and future studies should leverage longitudinal data to validate our developmental findings. The resting-state acquisitions were also different between the two datasets: while POND participants viewed a movie, HBN participants were instructed to fixate on a cross. Thus, the harmonization that was applied across both datasets may be inadvertently removing effects specific to each type of acquisition. We also only considered regional measures of cortical thickness and surface area, which greatly reduced the computational expenses of our analysis; future work could take a less limited spatial resolution and examine between-dataset differences in vertex-wise measures of brain structure. Finally, given the sample size required to establish reproducible brain-behaviour associations (Marek et al., 2022), the contribution of the identified factors to deviations from age models should be validated in larger samples.

The replicability crisis has posed a significant challenge to the neuroscience field and to the broader scientific community. In this context, we have questioned the extent to which age-related changes of brain structure and function are consistent between two independent cohorts of individuals with neurodevelopmental conditions. While deriving the neurobiological changes that occur during development is often a specific research goal, the presence or absence of replicable developmental effects is also highly relevant for all investigations which include age at any step in the methodology. Given our findings that age-related differences are not consistent across samples, this urges future consortium-based studies to carefully examine age-related changes within each dataset prior to applying the same assumptions (e.g., linearity) across datasets. This is particularly pertinent in the context of between-dataset differences in race/ethnicity and head motion, which may be important covariates given their observed associations with deviations from age-related models. While previous work has established cohort-specific effects in neurotypical samples, the heterogeneity associated with neurodivergence makes this finding specifically relevant to neurodivergent populations, where it is less likely that two samples have comparable characteristics. While inconsistencies were observed in both males and females, effect sizes were, on average, larger in the females compared to the males, suggesting that stratification approaches may be more suitable for addressing dataset-specific age-related changes in neurobiology.

## Data availability statement

Data is available from the Province of Ontario Neurodevelopmental Disorders (POND) network (exported April 2023; now available via a controlled data release through Ontario Brain Institute’s Brain-CODE: https://www.braincode.ca/) and the Healthy Brain Network (Release 9; http://fcon_1000.projects.nitrc.org/indi/cmi_healthy_brain_network/) datasets.

## Ethics approval statement

The current study was approved by the institution’s research ethics board, and both consortia studies were approved by the appropriate boards.

## Acknowledgements

We would like to thank all participants and their families for their involvement in this study. We would also like to thank all individuals who work for POND for their invaluable assistance in data collection.

This manuscript was prepared using a limited access dataset obtained from the Child Mind Institute Biobank, the Healthy Brain Network (HBN). This manuscript reflects the views of the authors and does not necessarily reflect the opinions or views of the Child Mind Institute. This research was conducted with the support of the Ontario Brain Institute (POND, PIs: Anagnostou/Lerch), an independent non-profit corporation, funded partially by the Ontario government. The opinions, results and conclusions are those of the authors and no endorsement by the Ontario Brain Institute is intended or should be inferred.

## Supplemental information

**Supplemental Table 1:** Acquisition parameters for the sMRI and rs-fMRI protocols for each site in the POND and HBN datasets.

| Dataset | Scanner | Scan type | Sequence | TR (ms) | TE (ms) | FA (°) | FOV (mm) | Voxel size (mm) | Scan time (min) |
| --- | --- | --- | --- | --- | --- | --- | --- | --- | --- |
| sMRI protocol |  |  |  |  |  |  |  |  |  |
| POND | SK: 3T TimTrio | – | MPRAGE | 2300 | 2.96 | 9 | 192×240×256 | 1 | 5.0 |
|  | QU: 3T TimTrio | – | MPRAGE | 2300 | 3.14 | 9 | 192×240×256 | 0.8 | 6.2 |
|  | SK: 3T Prisma<sup>fit</sup> | – | MPRAGE | 1870 | 3.14 | 9 | 192×240×256 | 0.8 | 5.0 |
|  | QU: 3T Prisma<sup>fit</sup> | – | MPRAGE | 1870 | 3.10 | 9 | 192×240×256 | 0.8 | 5.0 |
|  | HB: 3T Prisma<sup>fit</sup> | – | MPRAGE | 1870 | 3.10 | 9 | 192×240×256 | 0.8 | 5.0 |
| HBN | CBIC: 3T Prisma<sup>fit</sup> | – | MPRAGE | 2500 | 3.15 | 8 | 179×256×256 | 0.8 | 7.00 |
|  | CUNY: 3T Prisma<sup>fit</sup> | – | MPRAGE | 2500 | 3.15 | 8 | 179×256×256 | 0.8 | 7.00 |
|  | RU: 3T TimTrio | – | MPRAGE | 2500 | 3.15 | 8 | 179×256×256 | 0.8 | 7.00 |
|  | SI: 1.5T Avanto | – | MPRAGE | 2730 | 1.64 | 7 | 176×256×256 | 1 | 6.53 |
| fMRI protocol |  |  |  |  |  |  |  |  |  |
| POND | SK: 3T TimTrio | Movies | EPI | 2340 | 30 | 70 | 224×224×140 | 3.5 | 5.1 |
|  | QU: 3T TimTrio | Inscapes | EPI | 2340 | 30 | 70 | 224×224×140 | 3.5 | 5.1 |
|  | SK: 3T Prisma<sup>fit</sup> | Inscapes | EPI | 1500 | 30 | 70 | 222×222×150 | 3.0 | 5.1 |
|  | QU: 3T Prisma<sup>fit</sup> | Inscapes | EPI | 1500 | 30 | 70 | 222×222×150 | 3.0 | 5.1 |
|  | HB: 3T Prisma<sup>fit</sup> | Inscapes | EPI | 1500 | 30 | 70 | 222×222×150 | 3.0 | 5.1 |
| HBN | CBIC: 3T Prisma<sup>fit</sup> | Fixation | EPI | 800 | 30 | 31 | 202×202×144 | 2.4 | 2×5.1 |
|  | CUNY: 3T Prisma<sup>fit</sup> | Fixation | EPI | 800 | 30 | 31 | 202×202×144 | 2.4 | 2×5.1 |
|  | RU: 3T TimTrio | Fixation | EPI | 800 | 30 | 31 | 202×202×144 | 2.4 | 2×5.1 |
|  | SI: 1.5T Avanto | Fixation | EPI | 1450 | 40 | 55 | 195×195×135 | 2.5 | 10.3 |
|  | SK: 3T TimTrio | Movies | EPI | 2340 | 30 | 70 | 224×224×140 | 3.5 | 5.0 |
|  | QU: 3T TimTrio | Inscapes | EPI | 2340 | 30 | 70 | 224×224×140 | 3.5 | 5.0 |

sMRI: structural magnetic resonance imaging; rs-fMRI: resting-state functional magnetic resonance imaging; POND: Province of Ontario Neurodevelopmental network; HBN: Healthy Brain Network; SK: Hospital for Sick Children; QU: Queen's University; HB: Holland Bloorview Kids Rehabilitation Hospital; CBIC: CitiGroup Cornell Brain Imaging Center; CUNY: the City University of New York Advanced Science Research Center; RU: Rutgers University; SI: Staten Island; TR: repetition time; TE: echo time; FA: flip angle; FOV: field of view

**Supplemental Table 2:** Summary of the phenotypic measures of core and co-occurring differences for the diagnostic categories.

| Domain | Assessment | Score | Range | Interpretation<sup>1</sup> | Acronym |
| --- | --- | --- | --- | --- | --- |
| Social communication difficulties | Social Communication Questionnaire (Rutter et al., 2003) | Total score | 0-39 | More severe difficulties with social communication | SCQ |
| Repetitive behaviours | Repetitive Behaviours Scale – Revised (Bodfish et al., 2000) | Total score | 0-129 | More severe repetitive behaviours | RBS-R |
| Inattention | Strengths and Weaknesses of ADHD-symptoms and Normal Behaviour (Swanson et al., 2012) | Inattention subscale | 0-9 | More severe difficulties with inattention | SWAN-I |
| Hyperactivity | Strengths and Weaknesses of ADHD-symptoms and Normal Behaviour (Swanson et al., 2012) | Hyperactivity subscale | 0-9 | More severe difficulties with hyperactivity/impulsivity | SWAN-HI |
| Obsessive-compulsive traits | Toronto Obsessive-Compulsive Scale (Nelson et al., 2001) | Total score | -63-63 | More severe obsessive-compulsive traits | TOCS |
|  | Child Behaviour Checklist (Nelson et al., 2001) | Obsessive-compulsive subscale | 0-16 | More severe obsessive-compulsive traits | CBCL-OCS |
| Intelligence | Age-appropriate Wechsler or Stanford-Binet scales | Full-scale intelligence quotient | 40-160 | Higher intelligence | FSIQ |
| Internalizing problems | Child Behaviour Checklist (Achenbach & Ruffle, 2000) | Internalizing problems subscale | 40-75 | Worse internalizing problems | CBCL-I |
| Externalizing problems | Child Behaviour Checklist (Achenbach & Ruffle, 2000) | Externalizing problems subscale | 40-75 | Worse externalizing problems | CBCL-E |

<sup>1</sup>Interpretation: relative to higher scores

**Supplemental Table 3:**
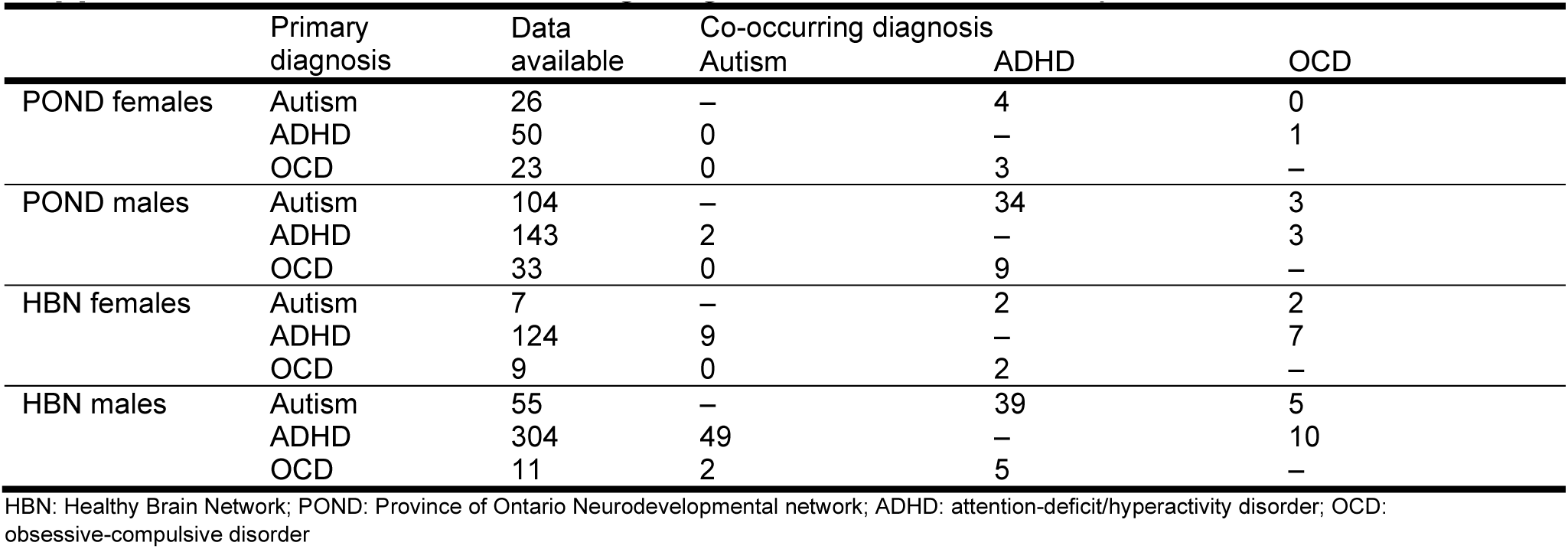
Co-occurring diagnosis in the sMRI sample.

|  | Primary diagnosis | Data available | Co-occurring diagnosis |  |  |
| --- | --- | --- | --- | --- | --- |
|  |  |  | Autism | ADHD | OCD |
| POND females | Autism | 26 | – | 4 | 0 |
|  | ADHD | 50 | 0 | – | 1 |
|  | OCD | 23 | 0 | 3 | – |
| POND males | Autism | 104 | – | 34 | 3 |
|  | ADHD | 143 | 2 | – | 3 |
|  | OCD | 33 | 0 | 9 | – |
| HBN females | Autism | 7 | – | 2 | 2 |
|  | ADHD | 124 | 9 | – | 7 |
|  | OCD | 9 | 0 | 2 | – |
| HBN males | Autism | 55 | – | 39 | 5 |
|  | ADHD | 304 | 49 | – | 10 |
|  | OCD | 11 | 2 | 5 | – |

HBN: Healthy Brain Network; POND: Province of Ontario Neurodevelopmental network; ADHD: attention-deficit/hyperactivity disorder; OCD: obsessive-compulsive disorder

**Supplemental Table 4:** Co-occurring diagnosis in the fMRI sample.

|  | Primary diagnosis | Data available | Co-occurring diagnosis |  |  |
| --- | --- | --- | --- | --- | --- |
|  |  |  | Autism | ADHD | OCD |
| POND females | Autism | 36 | – | 0 | 0 |
|  | ADHD | 37 | 0 | – | 1 |
|  | OCD | 30 | 0 | 8 | – |
| POND males | Autism | 114 | – | 49 | 4 |
|  | ADHD | 119 | 2 | – | 2 |
|  | OCD | 45 | 0 | 17 | – |
| HBN females | Autism | 7 | – | 4 | 2 |
|  | ADHD | 130 | 8 | – | 6 |
|  | OCD | 12 | 0 | 1 | – |
| HBN males | Autism | 53 | – | 35 | 5 |
|  | ADHD | 321 | 49 | – | 10 |
|  | OCD | 7 | 2 | 4 | – |

HBN: Healthy Brain Network; POND: Province of Ontario Neurodevelopmental network; ADHD: attention-deficit/hyperactivity disorder; OCD: obsessive-compulsive disorder

**Supplemental Figure 1:**
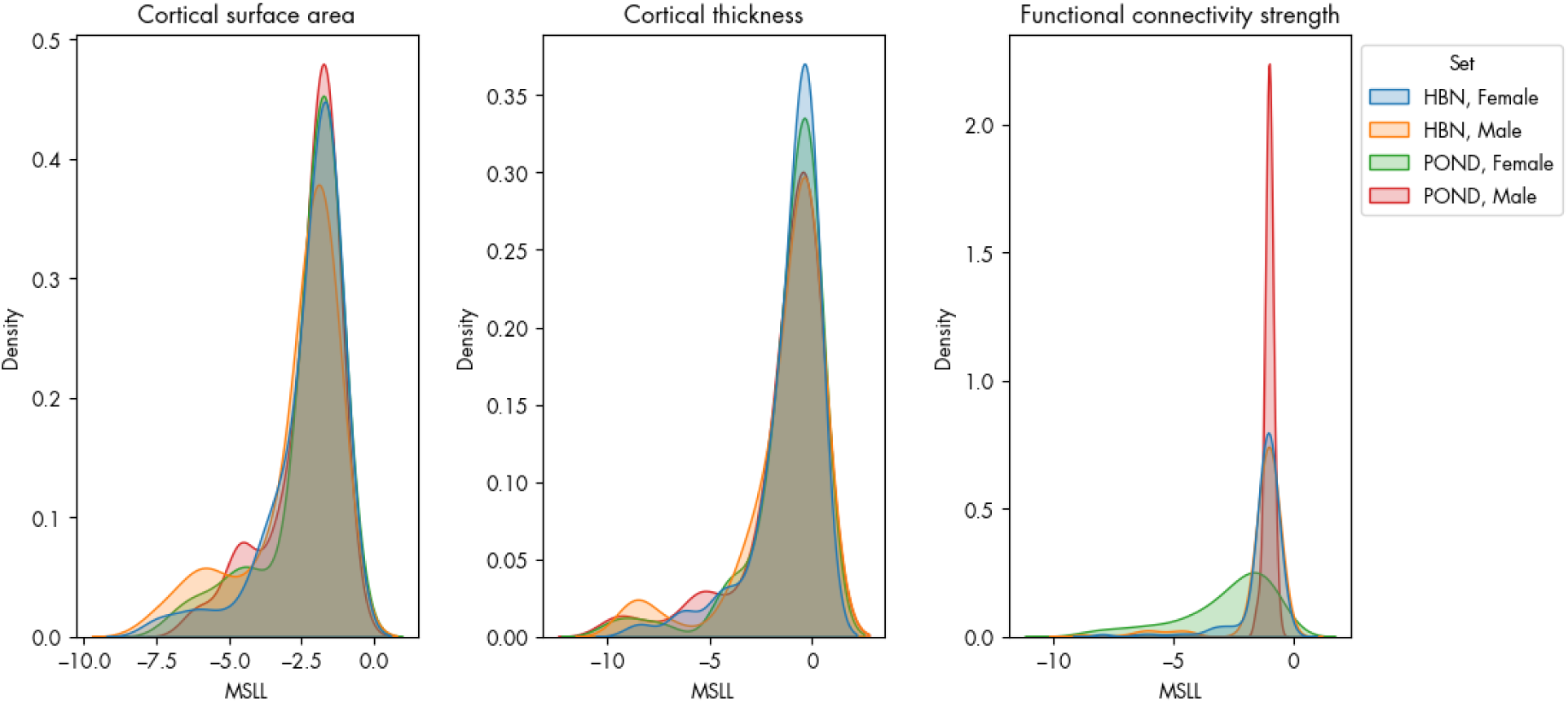
The distribution of mean squared log loss (MSLL) across all brain regions for the BLR models for cortical surface area, cortical thickness, and functional connectivity strength. MSLL was computed using data from the inner test set (HBN females, HBN males, POND females, and POND males).

**Supplemental Figure 2:**
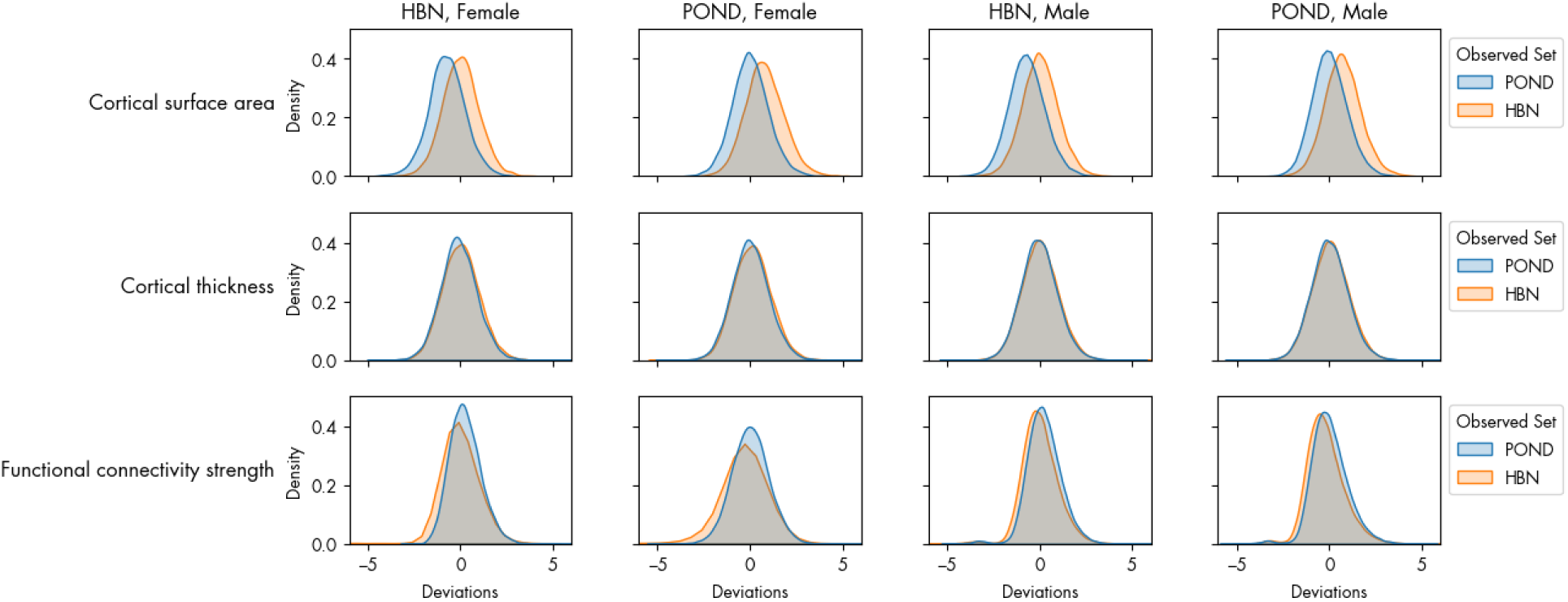
The distribution of deviation scores across all brain regions for the BLR models for cortical surface area, cortical thickness, and functional connectivity strength. Deviations were computed for the inner test set (HBN females, HBN males, POND females, and POND males) and its corresponding external test set.

**Supplemental Figure 3:**
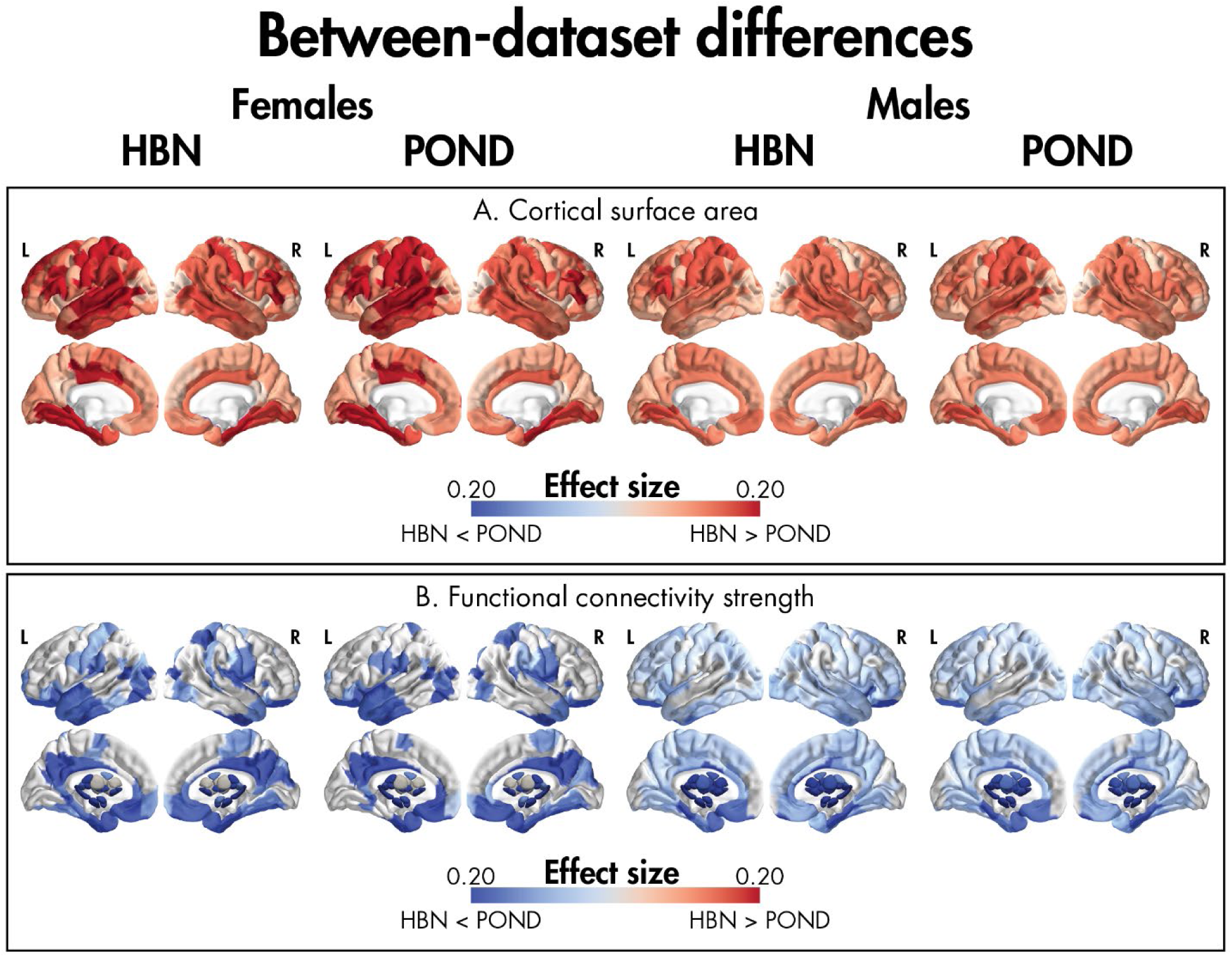
Effect sizes (eta-squared) for significant between-dataset differences in deviations from age-related models of cortical surface area (A) and functional connectivity strength (B) using a 70%/30% train/test split for the females (left) and males (right) when the models were built using each dataset (HBN: Healthy Brain Network, POND: Province of Ontario Neurodevelopmental network). No differences in cortical thickness were observed.

**Supplemental Figure 4:**
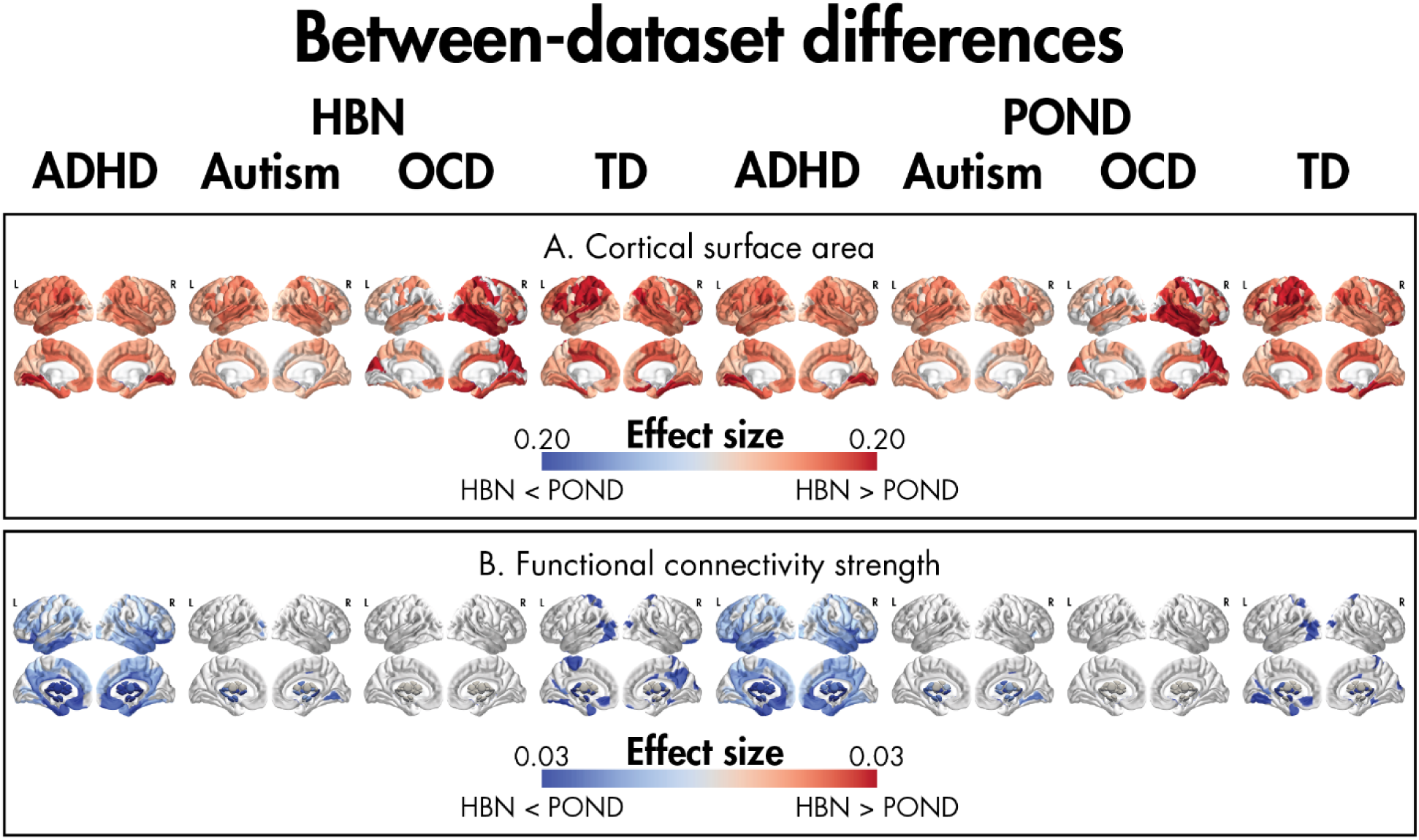
Effect sizes (eta-squared) for significant between-dataset differences in deviations from age-related models of cortical surface area (A) and functional connectivity strength (B) within each diagnosis and across sex when the models were built using each dataset (HBN: Healthy Brain Network, POND: Province of Ontario Neurodevelopmental network).

**Supplemental Figure 5:**
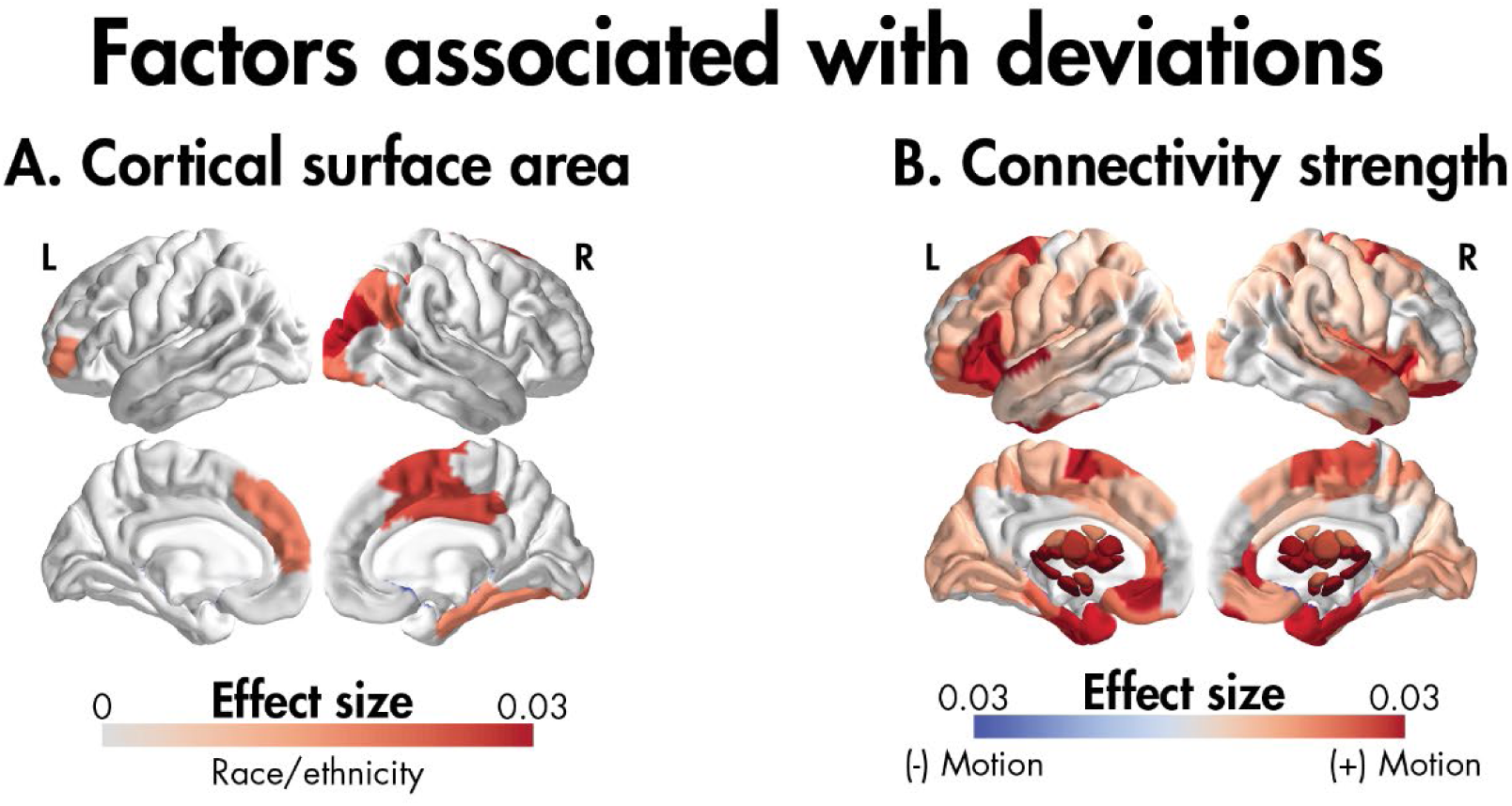
Significant association between (A) the deviations from age models of cortical surface area and race/ethnicity and (B) the deviations from connectivity strength models and head motion.

